## Supplemental figures and table for "Unraveling Enteroendocrine Cell lineage dynamics and associated gene regulatory networks during intestinal development"

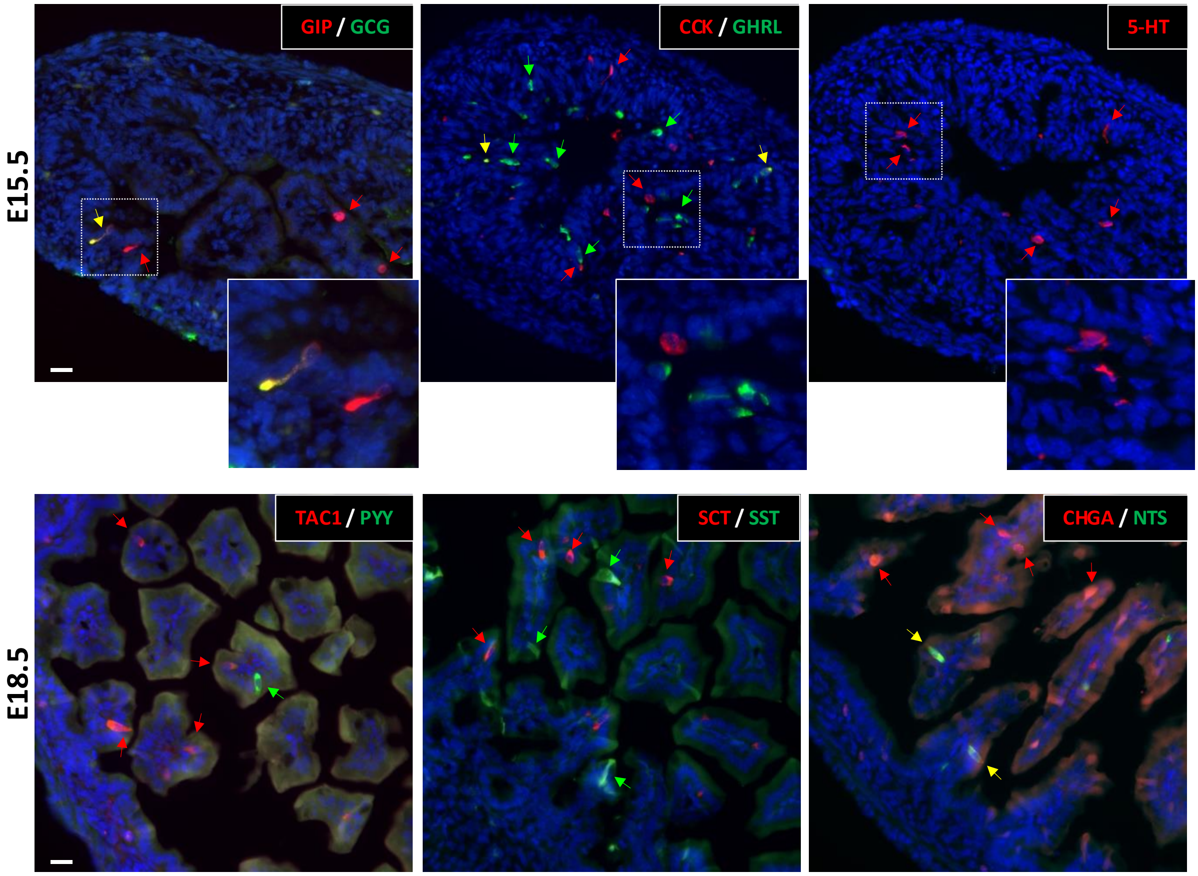


Sup. Figure 1

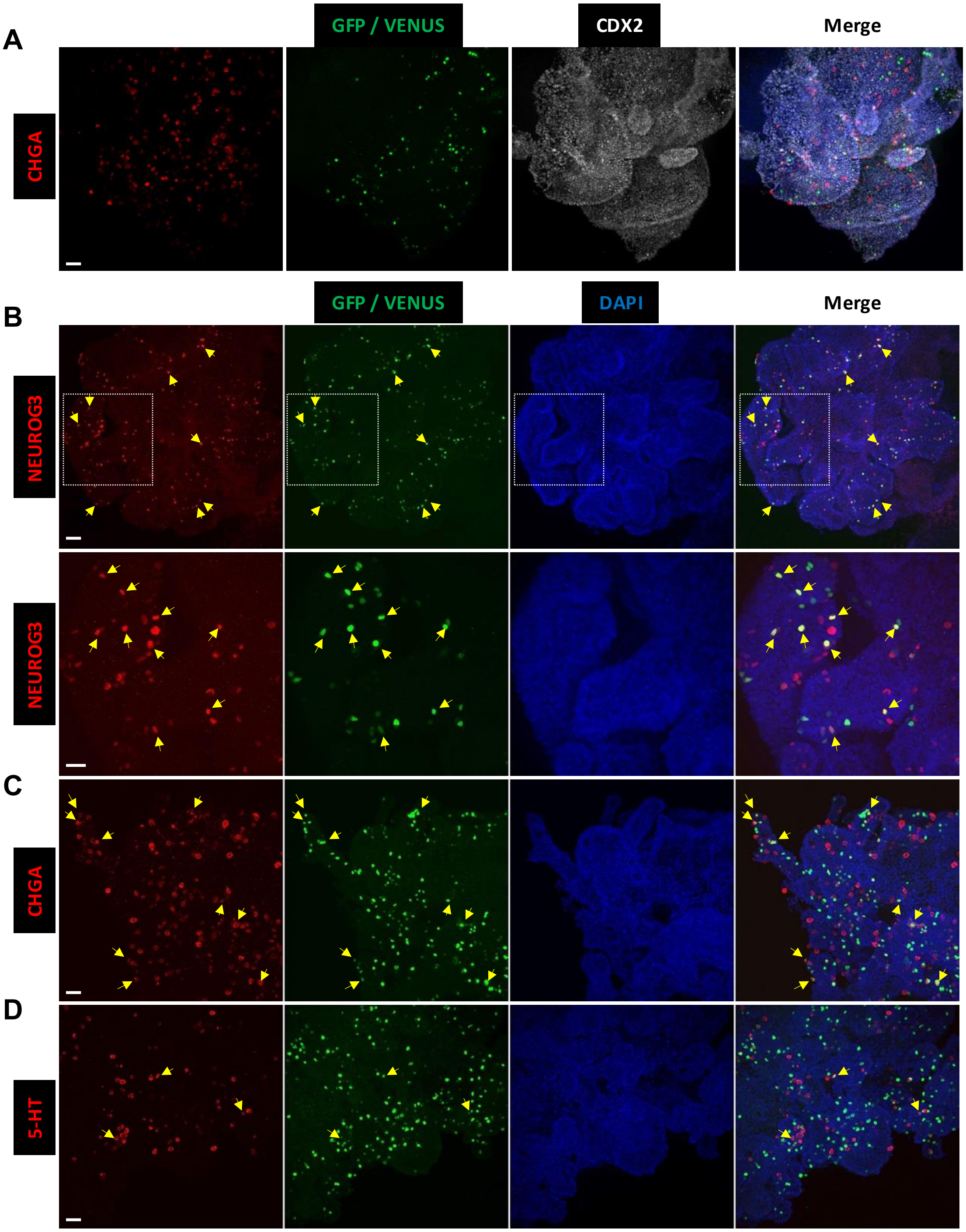


Supplementary Figure 2

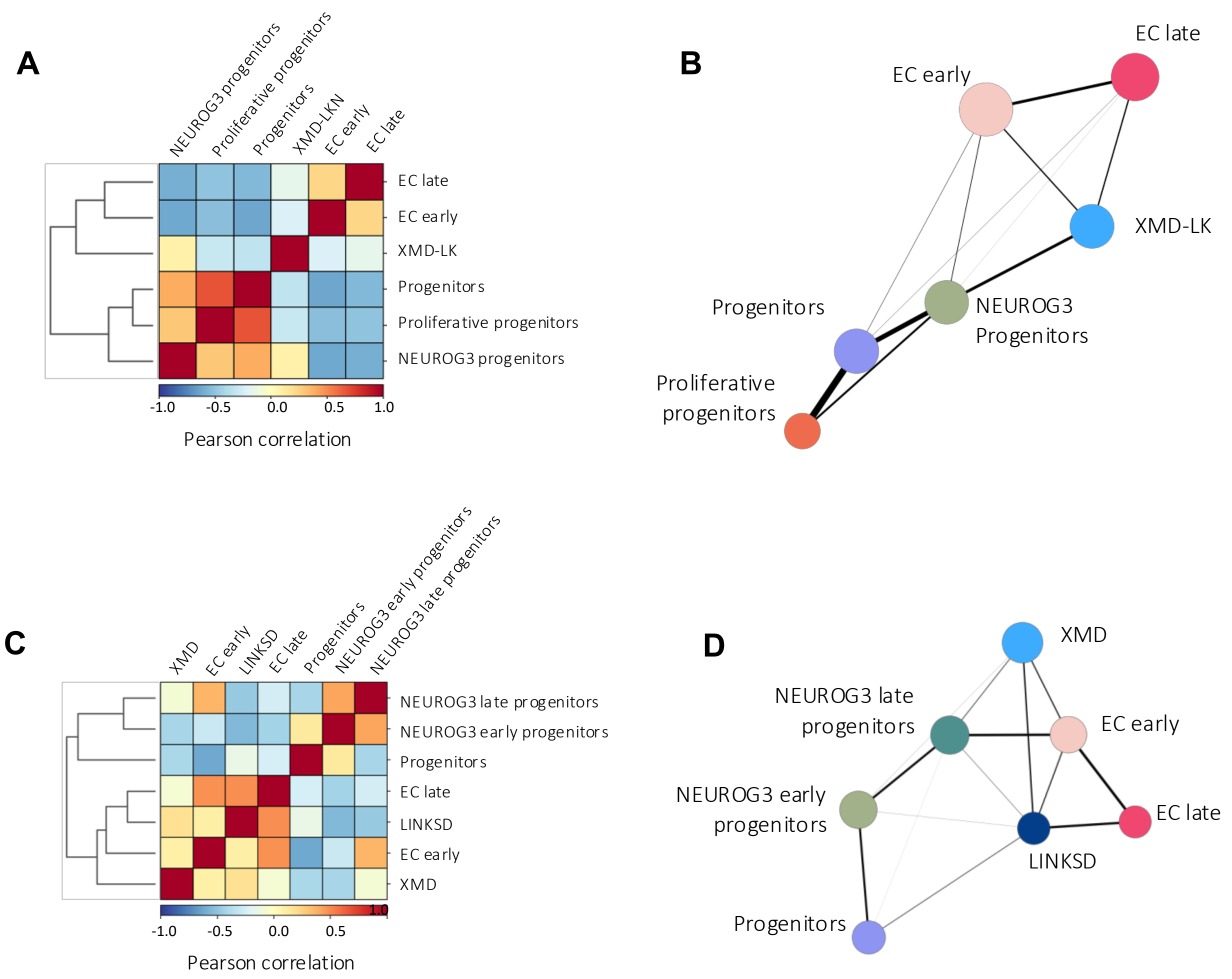


Supplementary figure 3

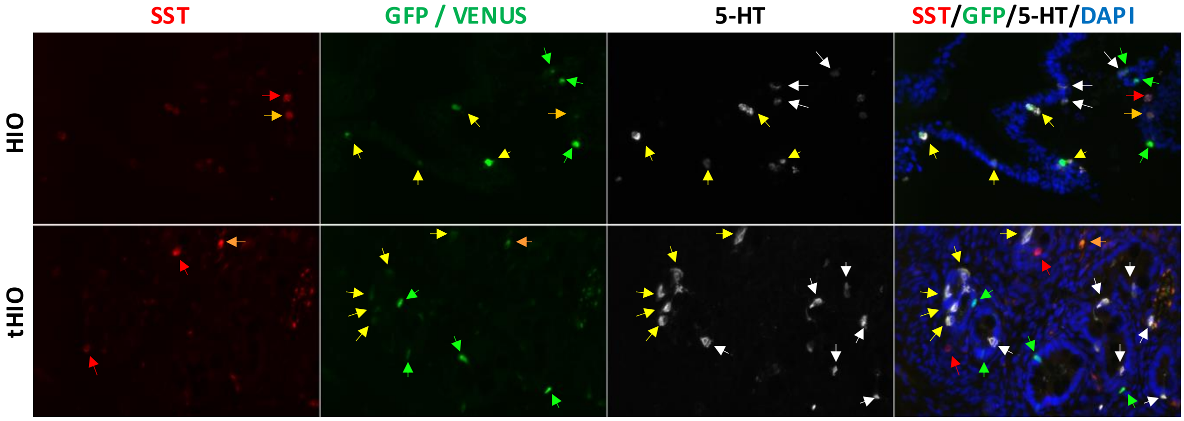


Supplementary figure 4

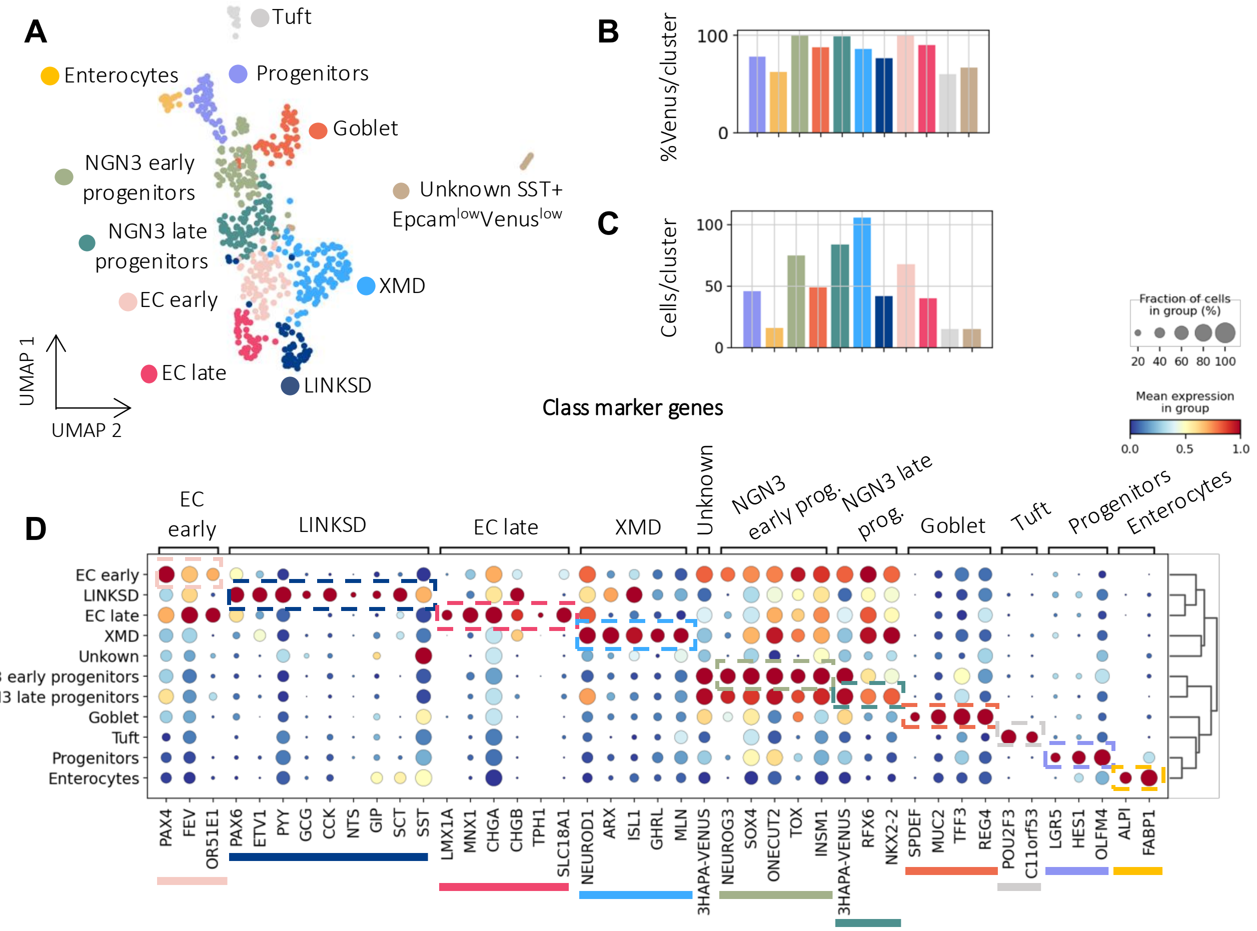


Supplementary figure 5

| Target | Host | Dilution | Reference | Manufacturer |
|---|---|---|---|---|
| NEUROG3 | Rabbit | 1/500 | IGBMC | IGBMC - G. Gradwohl |
| Ghrelin (GHRL) | Rabbit | 1/1000 | IGBMC | IGBMC - C. Tomasetto |
| Cholecystokinin (CCK) | Goat | 1/50 | Sc21617 | SantaCruz |
| GIP | Goat | 1/200 | Sc23554 | SantaCruz |
| Glucagon (GCG) | Guinea pig | 1/2000 | A-4031-01F | Linco |
| Serotonin (5-HT) | Goat | 1/2000 | Ab66047 | Abcam |
| Somatostatin (SST) | Rat | 1/1000 | MAB354 | Chemicon |
| Secretin (SCT) | Goat | 1/50 | Sc26630 | SantaCruz |
| GFP | Chicken | 1/2000 | ab13970 | Abcam |
| Chromogranin A (CHGA) | Goat | 1/500 | sc-1488 | SantaCruz |
| Tac1 (substance P) | Rat | 1/200 | MAB356 | Chemicon |
| Peptide YY (PYY) | Rabbit | 1/500 | H-059-03 | Phoenix |
| Neurotensin (NTS) | Rabbit | 1/500 | H-048-03 | Phoenix |

Table 1: List of primary antibodies used in the study
